# Integrating distal and proximal information to predict gene expression via a densely connected convolutional neural network

**DOI:** 10.1101/341214

**Authors:** Wanwen Zeng, Yong Wang, Rui Jiang

## Abstract

**Motivation:** Interactions among such cis-regulatory elements as enhancers and promoters are main driving forces shaping context-specific chromatin structure and gene expression. Although there have been computational methods for predicting gene expression from genomic and epigenomic information, most of them overlook long-range enhancer-promoter interactions, due to the difficulty in precisely linking regulatory enhancers to target genes. Recently, a novel high-throughput experimental approach named HiChIP has been developed and generating comprehensive data on high-resolution interactions between promoters and distal enhancers. On the other hand, plenty of studies have suggested that deep learning achieves state-of-the-art performance in epigenomic signal prediction, and thus promoting the understanding of regulatory elements. In consideration of these two factors, we integrate proximal promoter sequences and HiChIP distal enhancer-promoter interactions to accurately model gene expression.

**Results:** We propose DeepExpression, a densely connected convolutional neural network to predict gene expression using both promoter sequences and enhancer-promoter interactions. We demonstrate that our model consistently outperforms baseline methods not only in the classification of binary gene expression status but also in the regression of continuous gene expression levels, in both cross-validation experiments and cross-cell lines predictions. We show that sequential promoter information is more informative than experimental enhancer information while enhancer-promoter interactions are most beneficial from those within ±100 kbp around the TSS of a gene. We finally visualize motifs in both promoter and enhancer regions and show the match of identified sequence signatures and known motifs. We expect to see a wide spectrum of applications using HiChIP data in deciphering the mechanism of gene regulation.

**Availability:** DeepExpression is freely available at https://github.com/wanwenzeng/DeepExpression.

**Supplementary information:** Supplementary data are available at *Bioinformatics* online.

## 1 Introduction

Gene regulation, as one of the fundamental problems in biology, explains how different types of cells in the human body emerge from the identical information encoded by the genome (Ozbudak, *et al.*, 2002). The transcription of a gene is an extremely intricate process that requires a complex set of interactions among a myriad of proteins and DNA sequences (Maston, *et al.*, 2006). The regulation of this process is accomplished in large part by promoters and enhancers, which are DNA sequences containing multiple binding sites for a variety of transcription factors (TFs) (Yao, *et al.*, 2015). Enhancers can activate transcription independent of their location, distance or orientation with respect to the promoters of genes (Heinz, *et al.*, 2013). Therefore, ever since the emergence of high-throughput experiments for quantifying gene expression (Wang, *et al.*, 2009), computational biologists have long been interested in how well gene expression can be inferred by using TFs and regulatory elements (Rockman and Kruglyak, 2006), for deciphering the mechanism of gene regulation.

In computational studies of gene regulation (Lee and Young, 2013), various experimental data related to one-dimensional (1D) epigenomic signals, including TFs binding (Li, *et al.*, 2014), histones modification (Karlic, *et al.*, 2010) and chromatin accessibility (Duren, *et al.*, 2017), are taken as features to predict gene expression. These methods for predicting gene expression mainly fall into two categories. The first class of methods predict whether gene expression is high or low under a binary classification formulation. For example, DeepChrome (Singh, *et al.*, 2016) used five histone markers in promoter regions with a convolutional neural network (CNN) to predict gene expression status. The second class of methods infer continuous gene expression levels under a regression framework, and thus can provide quantitative predictions. For example, Ouyang *et al.* used ChIP-Seq data of 12 TFs in mESC cells with a linear regression model to predict gene expression (Ouyang, *et al.*, 2009). Karlić *et al.* collected nineteen histones modification in promoter regions in mouse embryonic stem cells and used a regression model for the prediction (Karlic, *et al.*, 2010). Dong *et al.* used twelve histone modification markers and chromatin accessibility in promoter regions with a two-step model for the prediction (Dong, *et al.*, 2012). They first constructed a random forest model to predict whether a promoter was active or not and then adopted a regression model to predict expression level of the corresponding gene. However, all these methods do not explicitly take enhancers and three-dimensional (3D) enhancer-promoter interactions into consideration thus far, probably due to both the difficulty in accurately linking enhancers to their target genes and the uncertainty of how strong these interactions will affect the gene expression (Mora, *et al.*, 2015).

The recent development of HiChIP (Mumbach, *et al.*, 2016), a high throughput experimental technique for sensitive and efficient analysis of protein-centric chromosome conformation, holds the promise to capture chromatin loops with high sensitivity and specificity. Compared with Hi-C (Belton, *et al.*, 2012) and chromatin interaction analysis by paired-end tag sequencing (ChIA-PET) (Li, *et al.*, 2014), HiChIP is able to measure protein-centric chromatin conformation in a rapid, efficient, technically simplified and high-resolution way. Among existing HiChIP studies, two of them stand out to show the ability of HiChIP in identifying enhancer-promoter interactions. First, Mumbach *et al.* evaluated H3K27ac, an enhancer-and promoter-associated mark, as a candidate factor to selectively interrogate enhancer-promoter interactions across the genome (Mumbach, *et al.*, 2017). Second, Weintraub *et al.* found the binding of YY1 to active enhancers and promoter-proximal elements and formed dimers that facilitated the interaction of these DNA elements (Weintraub, *et al.*, 2017). Therefore, HiChIP experiments of H3K27ac and YY1 have been developed to identify high-confidence 3D chromatin loops focused around enhancer-promoter interactions. These data sets provide valuable raw materials to study regulatory functions of enhancer-promoter interactions on gene expression.

Besides the rapid progress in biological experiments, recently, deep learning techniques have achieved the state-of-the-art performance on many bioinformatics applications such as regulatory site identification (Alipanahi, *et al.*, 2015) and biomedical image classification (Krizhevsky, *et al.*, 2017). A deep learning model automatically learns a complex non-linear function that maps inputs onto outputs, eliminating the need to use hand-crafted features or rules. As a representative model, CNN captures local characteristics in an input sample and learn important features that help make final predictions (LeCun, *et al.*, 2015). In the recent years, CNNs have been successfully used in a wide spectrum of fields such as computer vision (Razavian, *et al.*, 2014) and natural language processing (Vinyals, *et al.*, 2015). In bioinformatics, CNNs have been used to predict regulatory elements (Min, *et al.*, 2017), chromatin accessibility (Liu, *et al.*, 2018) and epigenetic states of a DNA fragment (Min, *et al.*, 2017; Zhou and Troyanskaya, 2015), as well as explain functional implications of genetic variants (Zhou and Troyanskaya, 2015).

Inspired by promising HiChIP experiments and advanced deep learning techniques, we introduce DeepExpression, a deep learning framework to model gene expression, with the consideration of enhancer, promoter, and their interactions. For distal enhancers, we adopt the state-of-the-art high-resolution 3D HiChIP experiments as features. For proximal promoters, we apply a recently developed deep learning model called densely connected convolution neural networks to extract epigenomic features in promoter regions. Validation experiments show that DeepExpression consistently outperforms several baseline methods not only in the classification of binary gene expression status but also in the regression of continuous gene expression levels. Model ablation analysis indicates that both the promoter information and enhancer information is informative for gene expression prediction. Furthermore, through a visualization strategy, we show that DeepExpression successfully captures sequence motifs in both promoter and enhancer regions, which are matched in the JASPAR database (Khan, *et al.*, 2018).

## 2 Methods

### 2.1 Data collection and preprocessing

We collected HiChIP data of H3K27ac for mouse embryonic stem cells (mESC) and identified corresponding RNA-seq data (Weintraub, *et al.*, 2017). We collected HiChIP data of YY1 for the HCT116, Jurkat and K562 cell lines (Weintraub, *et al.*, 2017) and identified corresponding RNA-seq data from the ENCODE project (Consortium, 2012). We extracted DNA fragments of 2,000 base pairs around transcription start site (TSS) of a gene as its promoter region. The summary of the data is shown in Table 1.

**Table 1.** Summary of data. Columns are the name of cell line, reference genome, number of genes, definition of promoters, and definition of enhancers in corresponding HiChIP experiment.

| Cell line | Species | Genes | Promoter | Enhancer |
| --- | --- | --- | --- | --- |
| mESC | mm9 | 18871 | TSS ±1000 bp | H3K27ac HiChIP |
| K562 | hg19 | 17899 | TSS ±1000 bp | YY1 HiChIP |
| Jurkat | hg19 | 17899 | TSS ±1000 bp | YY1 HiChIP |
| HCT116 | hg19 | 17899 | TSS ±1000 bp | YY1 HiChIP |

We followed the preprocessing pipeline described in (Weintraub, *et al.*, 2017) to deal with RNA-seq data. Taking mESC as an example, raw fasta data was aligned and quantified by using kallisto (version 0.43.0) (Bray, *et al.*, 2016) with the RefSeq transcriptome of mm9, resulting in estimated transcript counts. These counts were then summated across all isoforms of a gene to obtain gene-level counts. Finally, gene expression levels were calculated by applying a logarithmic transform of base 10 to gene-level counts, after adding a pseudocount of α (α = 0.1), and then quantile normalized across samples.

We followed the preprocessing pipeline described in (Weintraub, *et al.*, 2017) to deal with HiChIP data. We processed HiChIP data by first identifying reads with a restriction fragment junction, i.e., a site where ligation occurred. Reads containing a restriction fragment junction were trimmed such that the information from 5’ to the junction was kept. Reads without restriction fragment junctions were left untrimmed. The resulting reads were then mapped using bowtie (Langmead, *et al.*, 2009) against the mm9 genome or hg19 genome assembly. All unmapped or repetitively mapped reads were discarded in further analysis. We then utilized a statistical method named Origami (Weintraub, *et al.*, 2017) to identify high-confidence interactions that represent specific chromatin contacts, among all putative interactions. Here, specific chromatin contacts are defined as those that are detected with greater frequency than expected, given the linear genomic distance between two contacting regions. Finally, to adjust for different sequencing depths, we divide interaction counts *n_ij_* for interaction *j* in sample *i* by the total read count of the sample *Ni* and then scale the result by multiplying the minimum read count of all samples *N*. After this procedure, the raw count *n_ij_* for inteaction *j* of sample *i* was converted into a normalized read count 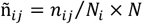.

Normalized interaction counts from a total of *n* replicate samples were averaged and then logarithmic transformed with base 2 after adding a pseudocount of 1 to characterize the interaction affinity of an interaction in a cell line. The value of HiChIP interaction signal is therefore

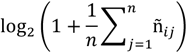

We divided the 2,000 kilo-basepair (kbp) DNA region (±1000 kbp) around the TSS of each gene into bins of length 5 kbp. Each bin includes adjacent positions of 5 kbp flanking the TSS of a gene. The value of HiChIP signals for a bin is then used as its input feature. These HiChIP signals represented long-range enhancer-promoter interactions.

### 2.2 Design of DeepExpression

As illustrated in Figure 1, DeepExpression consists of three modules. First, a proximal promoter module is used to extract features from DNA sequences in promoter regions. Second, a distal enhancer-promoter interaction module is used to extract features of HiChiP enhancer-promoter interactions signals. Finally, a joint module integrates outputs of the proximal promoters and distal enhancer-promoter interaction modules to produces a predicted gene expression signal.

**Figure 1.**
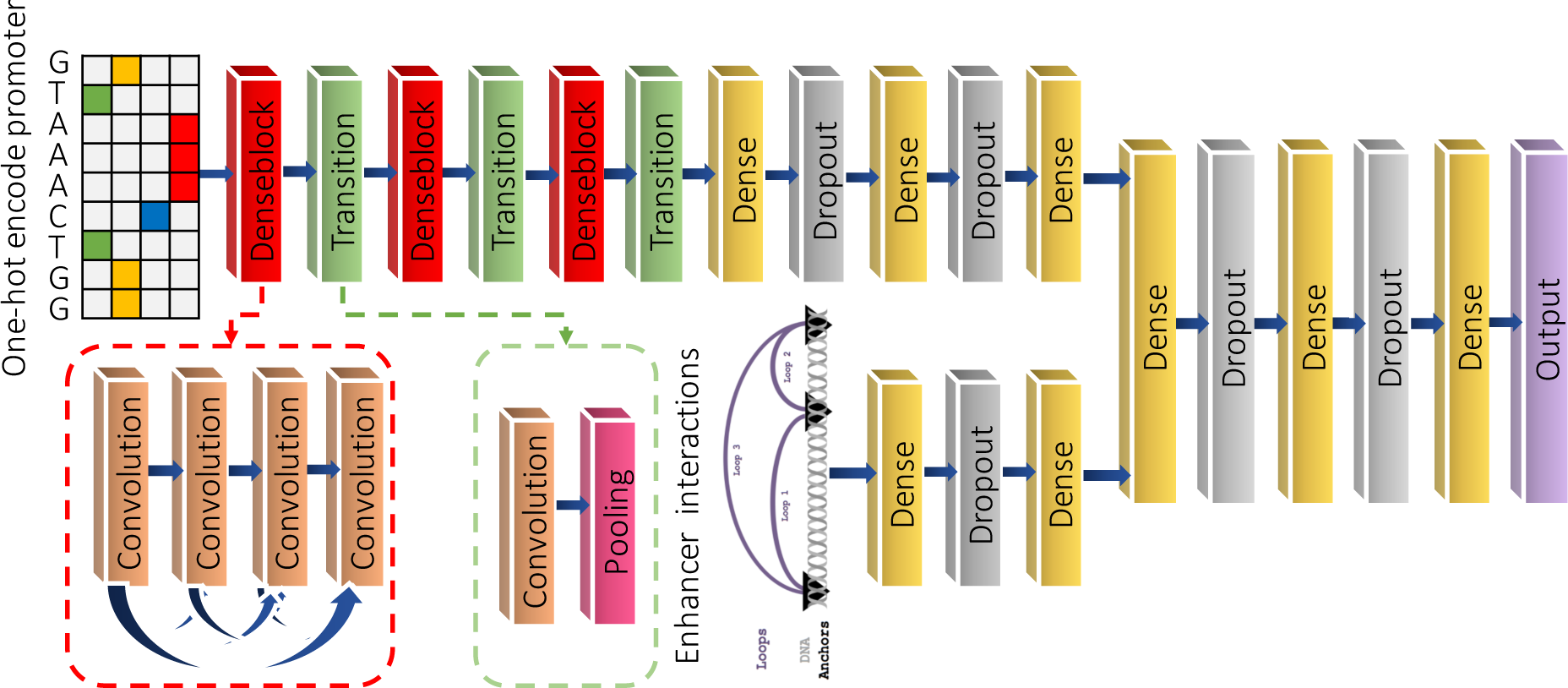
The graphical illustration of DeepExpression. First, a sequential promoter module is pre-trained to extract features from the input promoter regions. Second, an experimental HiChIP enhancer-promoter interactions module is adopted to fine-tune DeepExpression. Finally, a joint module integrates outputs of the promoter and enhancer modules to predict the gene expression.

**Figure 2.**
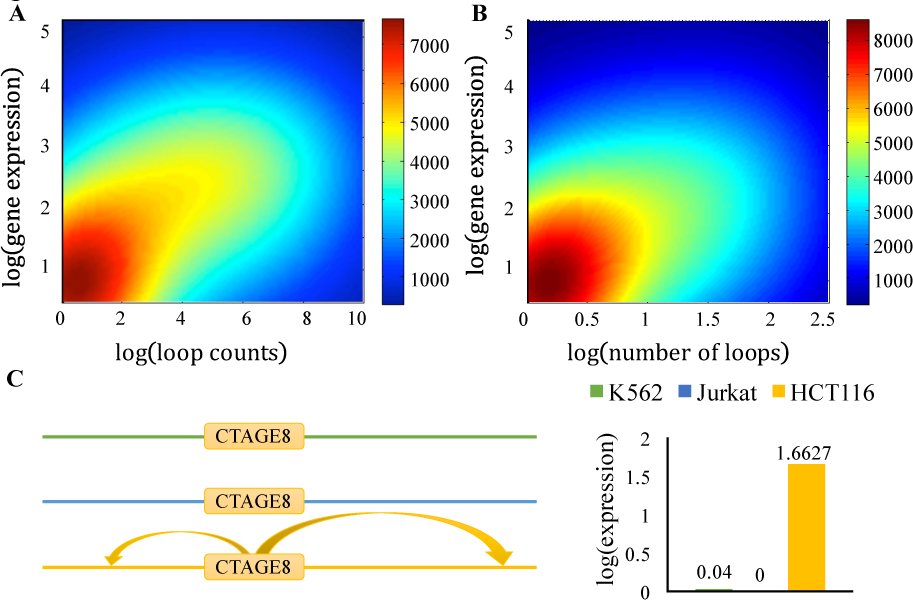
A) Scatter plot of HiChIP loop counts and gene expression (PCC: 0.623). The color bar on the right indicates the density of the scatter plot. B) Scatter plot of the number HiChIP loop counts and gene expression (PCC: 0.583). The color bar on the right indicates the density of the scatter plot. C) Visualization of the HiChIP loops of CTAG8, the expression of CTAG8 in K562, Jurkat and HCT116 cell lines respectively.

#### Proximal promoter module as a densely connected convolutional neural network

The proximal promoter module consists of three main components: a one-hot encoding input layer, three densely connected convolution blocks and three fully connected layers.

The one-hot encoding layer converts a DNA fragment into a numerical representation for downstream processing. It encodes the nucleotide in each position as a four-dimensional one-hot binary vector, in which each element represents one type of nucleotide: A, C, G, and T. The encoding layer then concatenates the binary vectors into a 4-by-2000 binary matrix, representing the whole 2000-bp target sequence.

The densely connected convolution blocks automatically extract features for an encoded DNA fragment. Recent advances in deep learning have shown that a classical convolutional neural network, though in general exhibiting high performance in prediction tasks, usually have hundreds of thousands of parameters involved, and thus often result in the severe overfitting problem on tasks with small training set sizes (Srivastava, *et al.*, 2014), like our data. To utilize parameters more efficiently and avoid the overfitting problem, a densely connected convolution network (Huang, *et al.*, 2017) connects all layers directly with each other, as schematically illustrated in Figure 1. In detail, in a block consists of *L* (*L* = 4) convolution layers, the input of the *l*-th layer is the concatenation of the feature-maps produced by all preceding layers 0,…,*l* −1, as

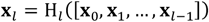

where H*_l_* denotes the concatenation operation. Meanwhile, the feature-map of the *l*-th layer are passed on to all *L* − *l* subsequent layers. This introduces *L*(*L* + 1)/2 connections in an *L*-layer network, instead of just *L*, as in a traditional architecture of convolutional neural networks. The convolution operation could be formulated as

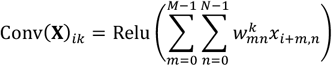

where **X** is the input matrix, *M* the size of the sliding window, *N* the number of input channels, 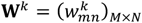 the weight matrix of the *k-*th convolution kernel with size *M* × *N*. In the first convolution layer, *N* is equal to 4. This first convolution process is equivalent to scanning a position weighted matrix (PWM) across the target sequence. In the other convolution layers, *N* is equal to the total number of convolutional kernels of all the preceding layers. The convolution layer then applies the rectified linear unit (ReLU) nonlinear function as

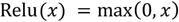

The pooling layer computes the maximum in each of the non-overlapping windows of size *M*, providing invariance to local shifts and reducing the number of parameters.

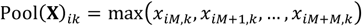

Three fully connected layers with 80, 40, and 40 units, respectively, performs linear transformations of the outputs of the previous layer, and applies the rectified linear unit nonlinear function. Multiple nonlinear layers enable the model to learn hierarchical representations of data with increasing levels of abstraction. Finally, the proximal promoter module transfers each sequential input to a 40-dimensional real vector.

#### Distal enhancer-promoter interaction module as a feedforward neural network

The distal enhancer-promoter interaction module receives 400-dimensional real vector HiChIP enhancer-promoter interactions signals as input. It uses two fully connected layers with 80 and 40 units to transform the one-dimensional numeric enhancer-promoter interaction strength input to a 40-dimensional real vector.

#### Joint expression prediction module as a feedforward neural network

The joint module integrates different features from both the proximal promoter and distal enhancer modules to predict gene expression. We merge outputs of these two modulus to form a feedforward neural network. For binary classification model, we use the softmax function to produce a probability output in the range of 0 to 1 that can easily and automatically be converted to output class values as 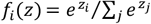, where *f_i_*(*z*) is the predicted probability that the input gene belongs to class *i* (i.e., +1 for high expression and −1 for low expression). For continuous regression model, we modified the output layer by replacing the softmax layer with a linear transformation layer.

### 2.3 Training of DeepExpression

We trained the proposed model in two main steps. First, in a model selection and pre-training step, we optimized the cross entropy loss function in the classification model and the mean squared error loss function in the regression model, using the RMSprop optimizer with a batch size of 4 and used dropout for the model regularization with a 0.5 dropout rate. We also applied the early stopping strategy (Erhan, et al., 2010) with the maximum number of iterations set to 60, and it would stop training after 5 epochs of unimproved loss on the validation set. We denote the model that has been trained up to this step as DeepExpression-seq.

Second, in a fine-tuning step, we incorporated the enhancer-promoter interaction module right before the joint output module. During the fine-tuning, we optimized the cross entropy or mean squared error loss function, only updating the weight parameters in the HiChIP enhancer-promoter interaction module. By fixing the weight parameters in the other layers, DeepExpression could avoid overfitting and effectively learn to incorporate the sequence representations with the HiChIP enhancer-promoter interactions information.

We adopted multi-fold cross-validation experiments to evaluate our model. Taking 10-fold as an example, we split each dataset into ten strictly non-overlapping groups by random sampling. In the validation, we used nine groups to train our model and the rest one as testing data. Data of the nine groups was further split as a training set and a validation set with ratio 0.8:0.1. The training set was used to adjust weights in the network, and the validation set was used to avoid overfitting. We implemented our method by using Keras, a deep learning library for Theano and Tensorflow. We used Theano as the backend, while the Tensorflow backend also generated very close results through our testing. The high-performance NVIDIA Tesla K80 GPU was used for model training.

### 2.4 Comparison with baseline machine learning models

To evaluate the performance of DeepExpression, we implemented three baseline methods for classification (logistic regression, SVM and random forest) and three methods for regression (linear regression, Lasso regression, and random forest regression). All the methods took both sequential and experimental data as input in accord with DeepExpression. For sequence data, we split the sequence of a DNA fragment into *k*-mers following the idea of gkmSVM (Ghandi, *et al.*, 2016). For experimental data, we took signals for bins to form an input vector. Baseline methods were implemented using the SciKit-learn library (http://scikit-learn.org).

We performed an internal 10-fold cross-validation experiment for model selection among regularization parameter and hyper-parameter configurations. For Lasso regression, we searched over 250 points that were evenly spaced between 10^−6^ and 10^6^ in log scale to optimize the regularization parameter. For SVM, we also searched over 250 points that were evenly spaced between 10^−6^ and 10^6^ in log scale to optimize the regularization parameter. For random forest classification and regression, we searched from all models selected from the following hyper-parameter configurations: the number of base estimators (chosen from [50, 100, 200, 300, 400, 500, 600, 700, 800, 900, 1000]) and the maximum depth of the individual regression estimators (chosen from [2, 4, 6, 8, 10]).

### 2.5 Statistical significance

We used two statistical tests in the SciPy library (http://scipy.org) for determining the statistical significance of prediction results. To determine the significance of the Pearson correlation between real and predicted gene expression levels, we used the two-tailed Student’s *t*-test with *n* − 2 degrees of freedom under the null hypothesis that the two sets of values are uncorrelated. For comparing the performance (measured by the AUC score for classification and Pearson correlation for regression) of two methods in cross validation experiments, we used one-sided Wilcoxon rank sum tests, which tests whether our method achieves higher performance than a baseline method.

### 2.6 Motif visualization

We convert each kernel of the first convolution layer into a position weight matrix (PWM) by scanning along input sequences for activated positions of the kernel and then calculating the PWM by pooling corresponding regions. We regard a position i as being activated if

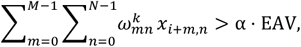

where *M* is the length of a kernel, *N* the number of input channels, α the control coefficient (0 < α < 1), and EAV the extreme activation value defined as

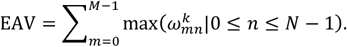

We set length of filters in the first convolutional layer to 8 and α to 0.7 in our visualization experiments. We identify putative motifs using the tool TomTom 4.11.2 (Gupta, *et al.*, 2007) with *E*-value threshold 0.05 to match PWMs identified by our method to the JASPAR database (Khan, *et al.*, 2018).

## 3 Results

### 3.1 HiChIP enhancer-promoter interactions are discriminative features for predicting gene expression

Since no previous studies have shown contributions of chromatin interactions to gene expression in a quantitative way, we first devised and tested the ability of HiChIP enhancer-promoter interactions to discriminate gene expression levels from the mESC cell line. We first simply summated all the loop counts of each gene and drew the heatmap of scatter plot of loop counts and gene expression. Figure 1A shows that the correlation between gene expression levels and total HiChIP loop counts is up to 0.623. We then counted the number of loops of each gene and drew the heatmap of scatter plot of the number of loops and gene expression. Figure 1B shows that correlation between gene expression and the number of HiChIP loop counts is also high, around 0.583. We could conclude that these HiChIP signals are highly correlated with gene expressions. Owing to the high-resolution and protein-centric HiChIP experiments, we could devise active enhancer-promoter interactions, which have huge impact on gene expression, from all the 3D interactions, and thus provides the promising features to predict gene expressions.

We further show a simple example of the relationship between gene expression and HiChIP signal in three human cell lines. CTAGE8 is an important paralog of CTAGE4, a gene associated with Cutaneous T Cell Lymphoma (Usener, *et al.*, 2003). We find that CTAGE4 only expresses in the HCT116 cell line and almost has no expression in K562 and Jurkat. From HiChIP data, no enhancer-CTAGE8 interactions are detected in K562 and Jurkat while two active enhancer-CTAGE8 interactions are found in HCT116.

From the above analysis, we could draw the conclusion that cell-type specific enhancer-promoter interactions obtained from HiChIP indeed provide useful regulatory information on gene expression, and thus are discriminative features for predicting gene expression.

### 3.2 DeepExpression accurately predicts binary gene expression status

We discretized the expression value of a gene to a binary status that indicates whether its expression is high or low. Given a specific cell line, we focused on all protein-coding genes collected from RefSeq, used the median of expression values across all such genes as a threshold, and assigned a positive label (+1) to a gene whose expression value is greater than or equal to the threshold and a negative label (-1) otherwise. We then systematically evaluated the performance of our method in capturing gene expression codes from the viewpoint of binary classification via a series of carefully designed multi-fold cross-validation experiments. We compared the performance of DeepExpression with three baseline methods, including logistic regression, SVM and random forest. We also proposed a variation of our model, named DeepExpression-seq, which discarded the HiChIP experimental data integration module and predicted gene expression using only DNA sequence information. We repeated the cross-validation experiments for different number of folds (5, 10, 15, 20), evaluated the performance of each method using a criterion called AUC, the area under the receiver operating characteristic curve (ROC), and reported the classification performance measured in AUC in Figure 3.

**Figure 3.**
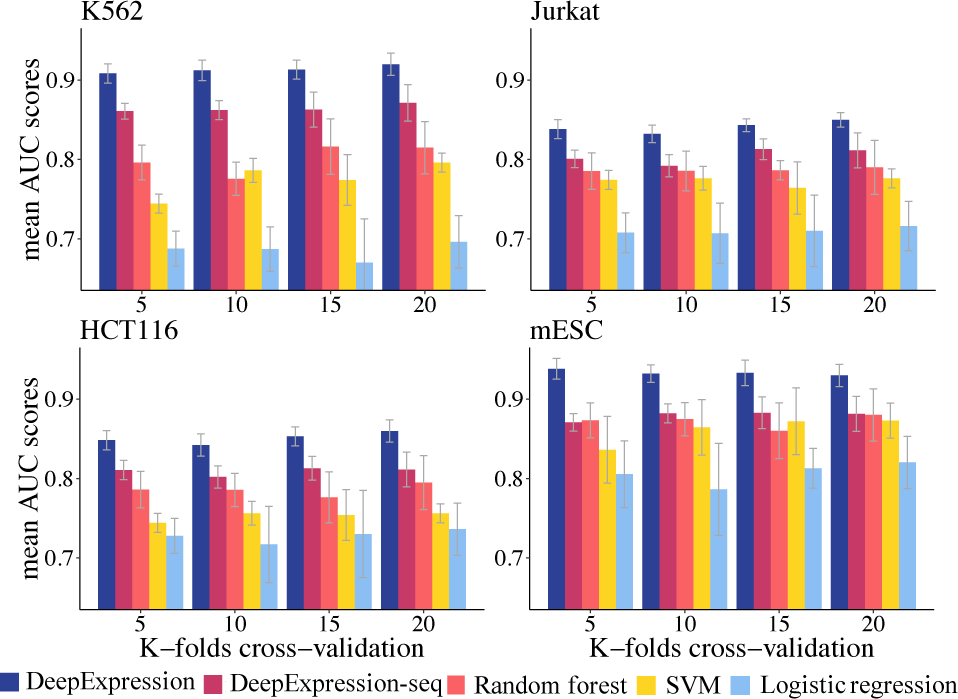
The classification performance measured in auROC at different *K*-folds cross validation experiments (*K*=5, 10, 15, 20).

As shown in the figure, our method consistently outperforms all the baseline methods. For example, in the 10-fold cross-validation experiment for mESC, the AUC scores of our method are 0.0574, 0.677 and 0.1458 higher than random forest, SVM and logistic regression, respectively. One-sided paired-sample Wilcoxon rank sum tests against the alternative hypothesis that our method achieves a higher AUC than a baseline method consistently show significant results (*p*-values < 3.1e-10 for all the three methods).

Our method also demonstrates much higher robustness than the baseline methods in Figure 3. For example, in the 10-fold cross-validation experiment for mESC, the variance of AUC scores of our method are 0.010, 0.024 and 0.047 lower than random forest, SVM and logistic regression, respectively. With such variances of AUCs calculated for cross-validation experiments of different folds for each cell line, one-sided paired-sample Wilcoxon rank sum tests against the alternative hypothesis that our method achieves smaller variance than a baseline method consistently show significant results (*p*-value < 6.1e-6 for all the three methods). We therefore conjecture that our method is not sensitive to the partition of training and test samples.

It is worth noting that the performance of DeepExpression-seq, which only uses DNA sequence information, is also superior to the three baseline methods and performs more steadily. For example, in the 10-fold cross-validation experiment for mESC, the AUCs of this model are 0.0073, 0.176 and 0.0957 higher than random forest, SVM and logistic regression, respectively. One-sided paired-sample Wilcoxon rank sum tests against the alternative hypothesis that our method achieves higher AUC scores than a baseline method consistently show significant results (*p*-values < 1.7e-4 for all the methods).

Finally, in terms of model training, benefit from the usage of dropout layers and the early stop strategy, the performance of DeepExpression on the test set is fairly close to that on the training set, indicating that our method is capable of avoiding overfitting. Taken together, these results suggest that deep learning outperforms the conventional machine learning methods for the prediction of gene expression based on target sequence composition and experimental enhancer-promoter interactions.

### 3.3 DeepExpression accurately regresses continuous gene expression levels

In the above classification experiments, we simply consider genes with two distinct status, namely highly or lowly expressed. However, degree of gene expression is actually a continuous variable, and hence binary classification models are unable to identify putative genes with different expression in more detailed level. To address this problem, we built a DeepExpression regression model to further recover the degree of gene expression levels. We modified the structure of the classification model by replacing the softmax output layer with a linear transformation layer. Besides, we used mean square error (MSE) as the loss function, thus forming the DeepExpression regression model.

Similar to the experiments in classification, we used our regression model to recover the gene expression with the same datasets. For evaluation purpose, we computed the Pearson Correlation Coefficient (PCC). We reported the regression performance measured in PCC in cross-validation experiments of different folds in Figure 4.

**Figure 4.**
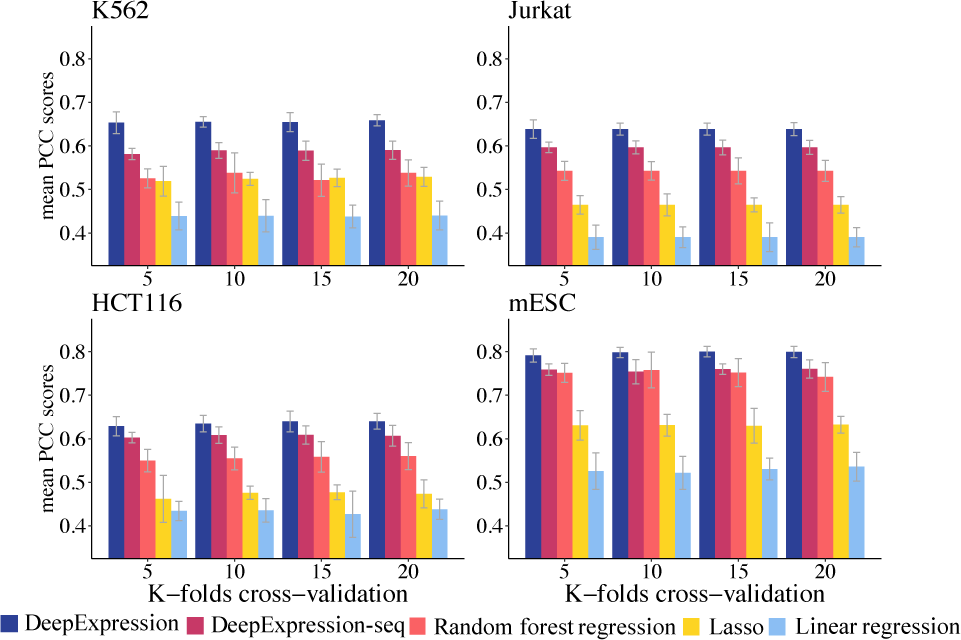
The regression performance measured in PCC at different *K*-folds cross validation experiments (*K*=5, 10, 15, 20).

As shown in the figure, our method consistently outperforms all the baseline methods. For example, in the 10-fold cross-validation experiment for mESC, the PCC of our method are on average 0.0404, 0.1669, and 0.2763 higher than random forest regression, Lasso and linear regression, respectively. One-sided paired-sample Wilcoxon rank sum tests against the alternative hypothesis that our method achieves higher PCCs than a baseline method consistently show significant results (*p*-values < 7.8e-13 for all the three methods). It is also worth noting that the performance of the DeepExpression-seq model is also superior to the three baseline methods and performs more steadily. For example, in the 10-fold cross-validation experiment for mESC, the PCC of DeepExpression-seq are on average 0.0020, 0.1227, and 0.2321 higher than random forest, Lasso and linear regression, respectively. One-sided paired-sample Wilcoxon rank sum tests against the alternative hypothesis that our method achieves higher PCCs than a baseline method consistently show significant results (*p*-values < 4.1e-5 for all the three methods).

Our method again demonstrates much higher robustness than the baseline methods in the regression task. With variances of PCCs calculated for cross-validation experiments of different folds for each cell line, one-sided paired-sample Wilcoxon rank sum tests as described in the previous section consistently suggest that our method achieves significantly smaller variance than a baseline method (*p*-value < 3.6e-8 for all the three methods).

Besides, we also noticed that all methods achieved higher performance in mESC than the other three cell lines in both classification and regression cases. We conjecture this is probably due to two reasons. First, gene regulation in stem cells is simpler than differentiated cells, and this reason may explain the situation that most gene expression prediction methods are only trained and predicted in mESC. Moreover, K562, Jurkat and HCT116 are all cancer cell lines, making it harder to model gene expression. Second, since H3K27ac has long been recognized as enhancers and promoters markers while YY1 is found to related to enhancer-promoter interaction recently, we presume that HiChIP experiments of H3K27ac might have higher accuracy than those of YY1 in capturing enhancer-promoter interactions.

In summary, the superior performance of our method in all cell lines indicate the general prediction ability of DeepExpression. With our regression model, we could determine and quantify the expression degree of a input gene with a continuous value. DeepExpression regression model hence provides us a broader way of predicting gene expression and inferring genes status.

### 3.4 Cross-cell line prediction

A HiChIP experiment provides a means of measuring how strong an enhancer regulates a target gene in a cell line. Since such regulatory relationships may be shared among cell lines, it could be possible to impute the expression of a gene in a cell line with the incorporation of HiChIP experimental data of other cell lines. To simulate this scenario, we performed a series of experiments for cross-cell line prediction by employing a collective scoring strategy. Given a cell line of interest and a gene, we used DeepExpression models trained on other cell lines to predict expression of the gene, and then averaged over such predictions to obtain the final prediction result of the gene for the given cell line.

We used the datasets of the three human cell lines to demonstrate the ability of our method to regress gene expression in a cross-cell line manner. For each of the three cell lines, we used the regression models of the other two cell lines to make predictions, and then averaged over the resulting two regression values to obtain a final regression value for a test gene. As shown in Figure 5, this cross-cell line predicting strategy is consistently superior to the three baseline methods. In more detail, the PCC of our method are on average 0.33, 0.30 and 0.27 higher than random forest, Lasso and linear regression, respectively.

**Figure 5.**
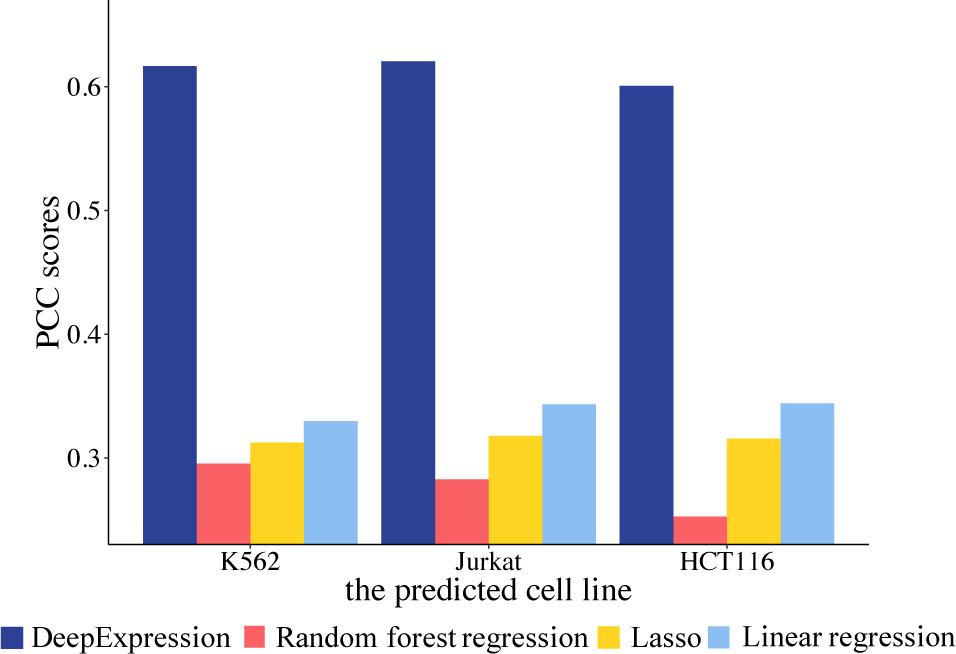
The cross-cell line performance measured in PCC at different independent testing cell lines.

These results suggest that by combining information of both promoters and enhancer-promoter interactions, we might learn the core code of gene regulation across different cell types. As the boosting of HiChIP enhancer-promoter interactions data, we expect to train DeepExpression in more and more cell lines, and consequently we could predict the gene expression for a new cell line that have not been studied yet. More importantly, we expect to learn the comprehensive and general gene regulation mechanisms with enhancer regulation across different cell lines.

### 3.5 Contributions of sequential and experimental features

The distal enhancer-promoter interaction module incorporates experimental HiChIP long-range enhancer-promoter interaction information into our methods. To prove that the experimental data is informative, we performed a model ablation analysis by repeating the cross-validation experiments with the enhancer-promoter interaction modules excluded. In a similar way, we excluded the promoter module to evaluate its contribution.

As shown in Figure 6, there are evident differences in the contributions of the proximal promoter and distal enhancer-promoter interaction module. Taking mESC as an example, after removing the promoter module, the mean PCCs decrease by 23.64%. When removing the enhancer-promoter module, however, the mean PCCs drop by 8.39%. Obviously, promoter sequences provide more information than enhancer-promoter interaction data to accurately predict gene expression. We speculate that there are two reasons accounting for the phenomenon. First, we incorporate HiChIP enhancer-promoter interactions using a simple fine-tuning way. These primitive feedforward networks might not catch all the information in HiChIP data. Second, since HiChIP is a newly developed experimental technique, there is no formal pipeline to process HiChIP data, and thus we might lose some information during the processing procedure. However, we could still conclude that using sequential promoter data and experimental enhancer jointly effectively improves the performance and both of them play important roles in predicting gene expression.

**Figure 6.**
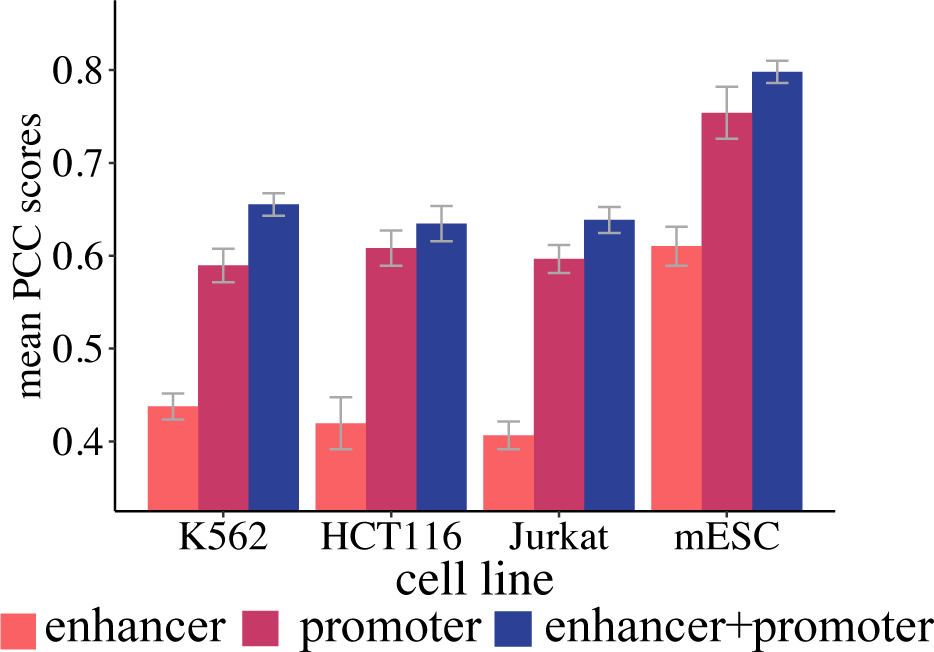
Contributions of sequential promoter and experimental enhancer features.

### 3.6 Contribution of enhancer-promoter interactions at different distances

To evaluate the contribution of enhancers at different distance, we carry out sensitivity analysis for different length of the HiChIP experimental input region. As shown in Figure 7, we find that when the length of the input region decreases, the performance of our model degrades slightly. For example, the mean PCC of DeepExpression on K562 is 0.6545 when the input region is ±1000 kbp around the TSS of each gene. Setting the input region to ±500 kbp around the TSS while retaining all the other parameters, we find the mean PCC is 0.6531, almost unchanged while the performance for input region ±100 kbp around the TSS drops a little bit to 0.6422. These results are consistent with our knowledge that most enhancers regulate the nearest genes (Spitz, 2016). From the above results, we could conclude that the information of HiChIP enhancer-promoter interactions is most beneficial from those within ±100 kbp around the TSS of a gene since there is no significant difference between performance under ±2000 kbp and ±100 kbp.

**Figure 7.**
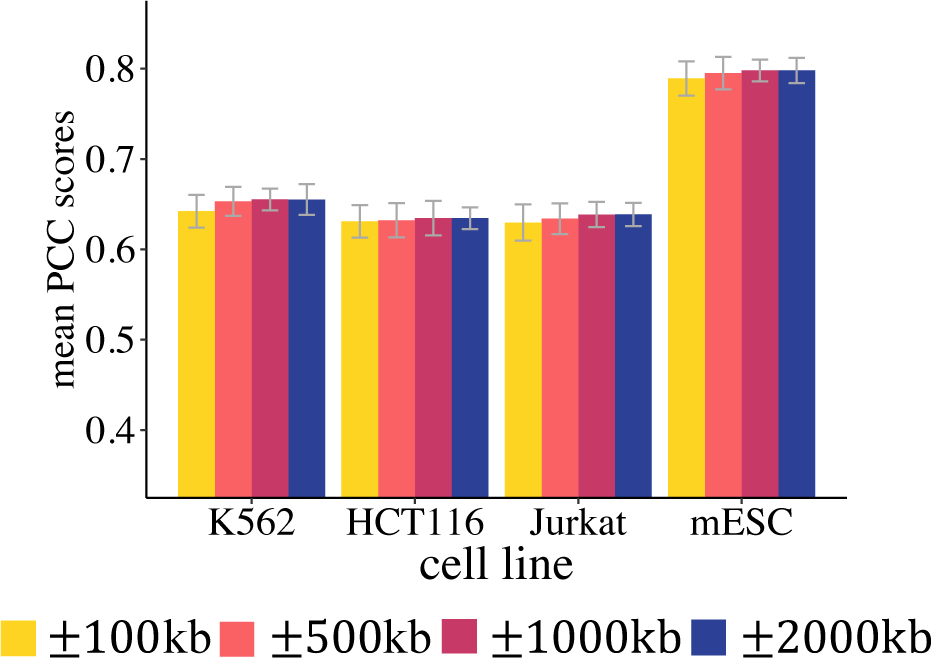
The mean PCC of different length of experimental input in each cell line.

Enhancers are notoriously difficult to locate and may reside at considerable distances from the transcription units on which they operate, for example, the enhancer of SHH is located more than 1Mb from its TSS. We considered that all genes can be divided into two group. Most genes are regulated by its neighboring enhancers (referred to as neighboring regulating genes), while some genes are regaled by long-range enhancers (referred to as long-range regulating genes). Since we do not have enough samples to distinguish long-range regulating genes from neighboring regulating genes now, we expect to carry out more detailed differential analysis between the long-range regulating genes and neighboring regulating genes when there are more data from HiChIP experiments in the near future.

### 3.7 DeepExpression recovers TF binding motifs in promoter regions

To demonstrate the interpretability of our model, we identified motifs learned from the first convolution layer of DeepExpression using the strategy detailed in Methods, and we compared these motifs with known Vertebrates motifs in the JASPAR database. Using motif comparison tool TomTom with significant *E*-value threshold 0.05, we matched about 65% (83/128) of motifs learned by DeepExpression to known motifs in different cell lines, as shown in Figure 8.

**Figure 8.**
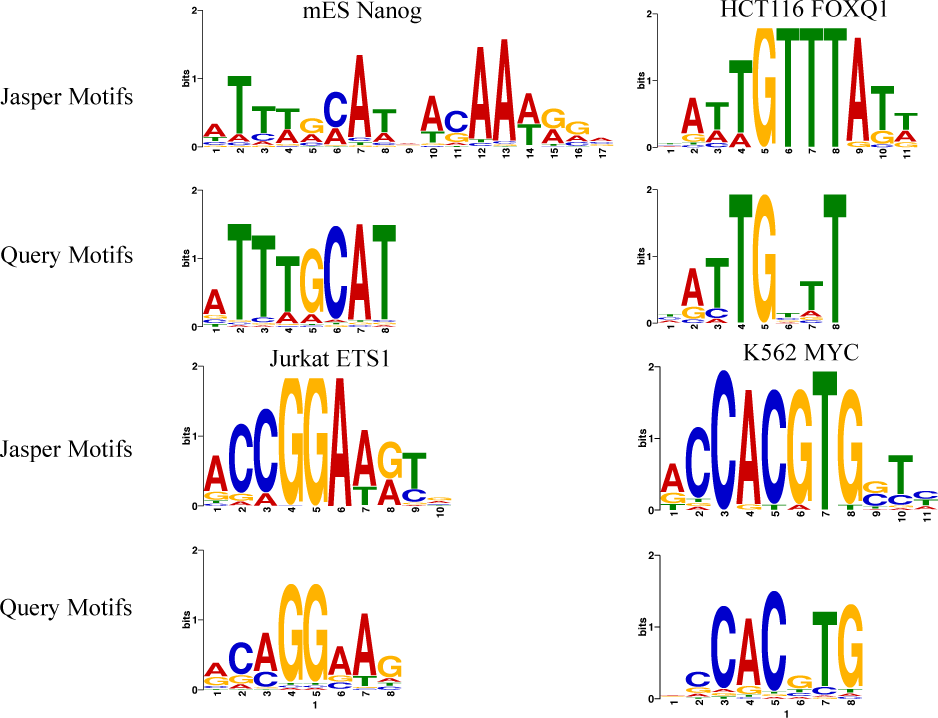
Motif visualization by promoter regions in each cell line.

To name a few, in mESC, DeepExpression recovers Nanog, a transcription factor involved in embryonic stem (ES) cell proliferation, renewal, and pluripotency. The protein encoded by this gene can block ES cell differentiation and can also autorepress its own expression in differentiating cells (Han, *et al.*, 2008). In HCT116, DeepExpression recovers FOXQ1, a member of the FOX gene family that is characterized by a conserved 110-amino acid DNA-binding motif called the forkhead or winged helix domain. FOX genes are involved in embryonic development, cell cycle regulation, tissue-specific gene expression, cell signaling, and tumorigenesis (Qiao, *et al.*, 2011). In Jurkat, DeepExpression discovers ETS1, which functions either as transcriptional activators or repressors of numerous genes and are involved in stem cell development, cell senescence and death, and tumorigenesis (Thomas, *et al.*, 1995). In K562, DeepExpression discovers MYC, a proto-oncogene and encodes a nuclear phosphoprotein that plays a role in cell cycle progression, apoptosis and cellular transformation (Gomez-Casares, *et al.*, 2013). The encoded protein forms a heterodimer with the related transcription factor MAX. This complex binds to the E box DNA consensus sequence and regulates the transcription of specific target genes. Amplification of this gene is frequently observed in numerous human cancers. Translocations involving this gene are associated with Burkitt lymphoma and multiple myeloma in human patients (Ceballos, *et al.*, 2000). To sum up, the powerful learning ability of DeepExpression could not only help us find potential TFs binding in specific cell line, but also guide us to find novel motifs which are not discovered by experiments yet.

### 3.8 DeepExpression recovers TF binding motifs in enhancer regions

To further make DeepExpression model more interpretable and convincing, we apply CisModule (Zhou and Wong, 2004) to visualize motifs learned from enhancer sequences in HiChIP data. CisModule is a hierarchical mixture *de novo* motif discovery algorithm, which develops a fully Bayesian approach for the simultaneous inference of TF modules and motif patterns based on their joint posterior distribution. We selected the highly interacted enhancer-promoter interactions in each cell line as the input and got the motif learned from these regions using CisModule. We then compared these motifs with known Vertebrates motifs in the JASPAR database. Using motif comparison tool TomTom with significant *E*-value threshold 0.05, we match about 92% (22/24) of motifs from HiChiP interactions. Moreover, the four distinguished motifs learned from promoter regions are also learned by the CisModule (Figure 9).

**Figure 9.**
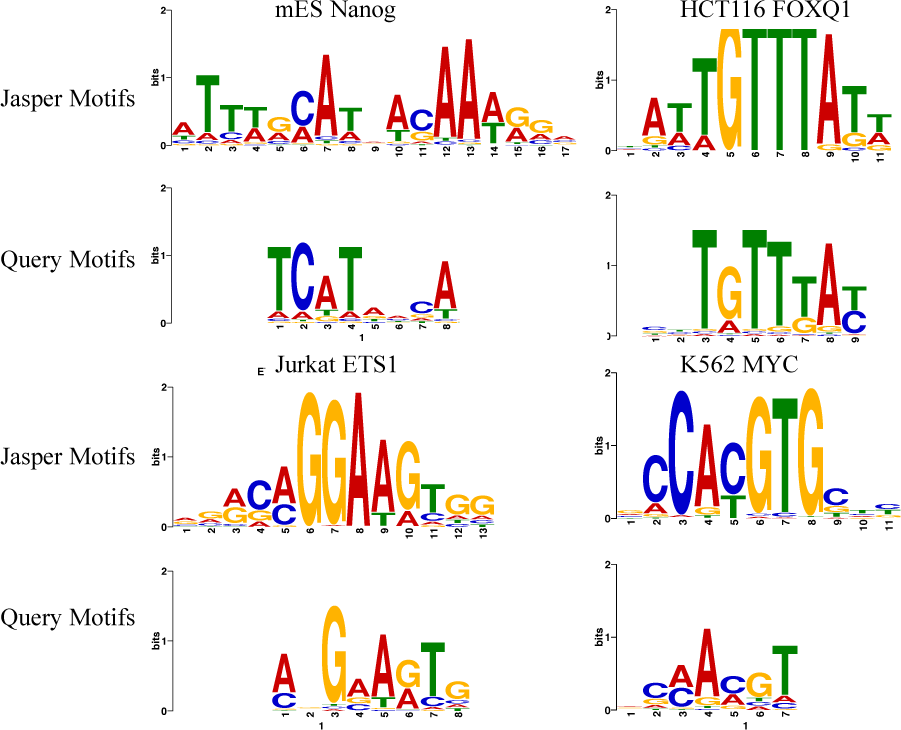
Motif visualization by enhancer regions in each cell line.

The consistency of TF discovered in promoter regions and enhancer regions demonstrate that the reason that jointly using both promoters and enhancers features could improve DeepExpression performance. Furthermore, the motif relevance in promoter regions and enhancer regions will be modelled explicitly in future version of DeepExpression.

## 4 Discussion

We have introduced a deep learning framework named DeepExpression to integrate DNA sequence information and enhancer-promoter interaction data for modelling gene expression. Through comprehensive validation experiments, we have shown that DeepExpression is superior to baseline methods in different cell lines and different species, capable of making cross-cell line predictions, and interpretable in extracted features.

DeepExpression is distinct from other methods for predicting gene expression in the following aspects. First, we adopt novel stat-of-the-art 3D HiChIP experimental features while existing methods only use 1D features like histone modification and chromatin accessibility (Shu, *et al.*, 2011). HiChIP defines the high-resolution landscape of enhancer-promoter regulation. Many complex features of the 3D enhancer connectome cannot simply be predicted from 1D data, demonstrating that it is necessary to employ these features. Second, we combine promoters and enhancer features together to model gene expression. Enhancers and promoters are the most important cis-regulatory elements and have huge impact on gene expression. Taking these two types of features into account, we could better model gene expression. Third, we apply densely connected convolution neural networks in DeepExpression. By reusing the features in each layer, densely connected convolution neural networks could use much less parameters and avoid overfitting in small training sets.

Nevertheless, our work can be further improved in several aspects. First, the incorporation of the long short-term memory (LSTM) network, a kind of recurrent neural network architectures, into our densely network framework may further improve the performance, because LSTM may be able to capture very long-range interaction in the sequence. In addition, the adaptation of an embedding representation of DNA sequences instead of the use of the one-hot encoding may also benefit the prediction accuracy (Min, *et al.*, 2017). Second, since we have shown that the first convolutional layer could capture motif information, researchers may use our model to learn the complex grammar of TF binding in specific cell lines. In addition, one can also explore interactions of motifs in higher convolutional layers. Third, our deep learning framework can possibly be adapted for the integration of other 3D functional elements interactions in the genome, including but not limited to silencers (Kolovos, *et al.*, 2012), repressors and insulators (Raab and Kamakaka, 2010). Forth, we should better model the motif information located in promoter and enhancer regions. Through section 3.6 and section 3.7, we could *de novo* discover import motifs in promoters and enhancers respectively. We could combine these motifs information in a unified framework to better model gene expression. Fifth, we could modify enhancer-promoter integration module in a more complicated way to better extract information from HiChIP data. Sixth, we expect to carry out cross-species predictions in our DeepExpression framework. We will integrate cross-species prediction once there are the same cell line in different species such as hESC and mESC. We look forward to decipher the enhancer-promoter interactions regulatory mechanism across species. Finally, our framework can be generalized for the prediction of functional impacts of genomic mutations and the prioritization of candidate variants in whole genome sequencing studies, thereby facilitating both research and practice of precision medicine (Alipanahi, *et al.*, 2015).

## Acknowledgements

We acknowledge the authors of YY1 Is a Structural Regulator of Enhancer-Promoter Loops, who provide us valuable data. Rui Jiang is a RONG professor at the Institute for Data Science, Tsinghua University.

